# The role of serotonin in modulating social competence in a cooperatively breeding fish

**DOI:** 10.1101/2023.07.18.549528

**Authors:** Diogo F. Antunes, Pia R. Stettler, Barbara Taborsky

## Abstract

Behavioural interactions between conspecifics rely on the appreciation of social cues, which is achieved through biochemical switching of pre-existing neurophysiological pathways. Serotonin is one of the major neurotransmitters in the central nervous system responsible for the modulation of physiological and behavioural traits, in particular social behaviour. The importance of serotonin for the ability to optimise ones social behaviour depending on available social information, that is, social competence, is yet unknown. Here we investigate how serotonin and the serotonin 1A receptor (5-HT_1A_) modulate social competence in a competitive context. In the cooperatively breeding cichlid *Neolamprologus pulcher*, we pharmacologically manipulated the serotonin availability and 5-HT_1A_ activity to test their effects on social behaviours during an asymmetric contest between the owner of a defended territory containing shelter and an intruder devoid of a territory. In this contest, the adequate response by the intruders, the focal individuals in our study, is to show submissive behaviour in order to avoid eviction from the vicinity of the shelter. While the serotonin enhancer Fluoxetine did not affect the frequency of submission towards territory owners, reducing serotonin by a low dosage of 4-Chloro-DL-phenylalanine (PCPA) increased submissive behaviour. Furthermore, threat displays towards territory owners were reduced at high dosages of Fluoxetine and also at the lowest dosage of PCPA. 5-HT_1A_ activation increased threat displays by intruders, indicating that this receptor may not be involved in regulating social competence. We conclude that serotonin, but not its receptor 5-HT_1A_ plays an important role in the regulation of social competence.

## 2. Introduction

Animal social behaviour is expressed in a range of different contexts, including courtship, resource competition, hierarchy formation and cooperation. Being able to show adequate social behaviours towards conspecifics and to adjust behaviour quickly to changing context has important fitness consequences [1,2]. The ability of individuals to optimize their social behaviour according to available social information is referred to as social competence [2]. For instance, in social species living in stable social hierarchies, adjusting the behaviour in response to social cues emitted by a multitude of different group members is important to maintain rank and group membership [3,4]. While the ultimate causation of animal social competence is well understood [2], the underlying proximate neurophysiological mechanisms of the ability to adjust social behaviours flexibly during ecologically relevant contexts remains largely unknown. The aim of this study is to understand the internal processes that are converting a sensory input into a behavioural output in a social fish species, in which the functional aspects of social behaviour are well understood (see reviews in 9,10). We focus on the role of serotonin (5-HT), a key neurotransmitter in the central nervous system, and one of its main receptors, 5-HT1A for the modulation of social competence in an ecologically relevant context.

Serotonin affects many behavioural and physiological functions such as aggression, locomotion, feeding, fear and stress (reviewed in 11–13). For instance, 5-HT influences the social status of an individual with more 5-HT inducing more subordinate behaviour both in lizards and fish [10,11]. Moreover, 5-HT reduces aggression in several invertebrate and vertebrate species (e.g. lobster: [12]; lizard: [13]; fish: [14]; mouse: [15]). For instance, during and after aggressive encounters, the 5-HT concentration in the prefrontal cortex of male Long-Evans rats decreased [16]. The pharmacological depletion of 5-HT by PCPA (4-Chloro-DL-phenylalanine), a 5-HT synthesis inhibitor, increased threat displays in rats [17]. The manipulation of the 5-HT system in male crickets (*Gryllus bimaculatus*) showed that 5-HT synthesis inhibitor leads to crickets to be more resilient to chronic defeat, as crickets with less 5-HT regain their aggressiveness after several consecutive defeats [18].

While 5-HT is an important mediator of aggressive interaction, its action is dependent on the activity of 5-HT receptors. When released to the synaptic cleft serotonin binds to different receptors that are localized mostly post-synaptically. Of these the receptors 1A, 2A, and 7 have been previously shown to influence sociality or mood [14,19–22]. 5-HT_1A_ is characterised by its high affinity for serotonin [23], and it plays an important role in modulating social behaviours [14,19,22,24]. Activation of 5-HT_1A_ leads to the inhibition of the adenylyl cyclase activity and opens potassium channels, causing neuronal hyperpolarization and a reduced firing rate [25]. 5-HT_1A_ occurs both as presynaptic autoreceptor and as postsynaptic heteroreceptor [26]. Activation of the autoreceptors leads to a feedback loop which down-regulates the serotonin release [27]. Pharmacological manipulations of 5-HT_1A_ can modulate aggressive behaviour. The application of a receptor agonist decreased aggression in a competitive context [14,19], whereas it increased aggression in members of a social group with stable hierarchy [22]. Besides its involvement in modulating aggressive interactions, 5-HT_1A_ also mediates socio-positive behaviours. In titi monkeys (*Callicebus cupreus,* [24]) and cichlid fish (*Neolamprologus pulcher*, [22]), injection a 5-HT_1A_ agonist reduced affiliative behaviour. In cleaner wrasses (*Labroides dimidiatus*), blocking the 5-HT_1A_ receptors reduced their motivation to engage in cleaning interactions, while the agonist of 5-HT_1A_ increased their motivation [19]. Additionally, 5-HT_1A_ also plays a role in the modulation of glucocorticoid stress responses (reviewed in: [9,28]).

Thus, while the importance of the serotonin system for modulating social behaviour in animals is unequivocal, its effects on regulating social interactions can be highly context dependent (e.g., see [14,19] vs. [22]). Here we studied the role of serotonin and its receptor 5-HT_1A_ in the regulation of social behaviour and social competence during contests over a resource between individuals of the cooperatively-breeding cichlid *N. pulcher*. We were particularly interested in the role of serotonin for the expression of submissive behaviour during contests, which has been rarely explored yet (but see [10,11]). We performed pharmacological manipulations of the serotonergic pathway by modifying the serotonin availability and the activity of 5-HT_1A_, by either increasing or decreasing serotonin availability, and increasing or decreasing the activity of 5-HT_1A_. This approach allowed us to disentangle between the general role of serotonin and the specific role of 5-HT_1A_ in modulating social interactions.

After drug application, we exposed the focal fish to an asymmetric contest over a resource. In this set-up, a focal fish is experimentally assigned to the role of an intruder during a territorial contest with a size-matched conspecific which owns the territory [29]. *N. pulcher* live in stable groups structured by size-based linear hierarchies [30]. These fish exhibit a large repertoire of aggressive, submissive and affiliative displays to maintain this hierarchy [6]. Their sociality has evolved as a strategy to cope with very high predation pressures [31,32]. Hence, access to a territory containing shelters to hide from predators is crucial for their survival [6]. In these fish, the 5-HT_1A_ modulates aggressive, submissive and affiliative interactions within social groups [22]. In the ‘asymmetric competition test’ used in our experiment, focal individuals do not possess an own shelter, which puts them in a disadvantage compared to a dominant territory owner, which monopolizes access to a shelter. In this test, the focal ‘intruder’ typically strives for tolerance by the dominant territory owner allowing them to remain in the vicinity of the shelter [29]. To achieve tolerance, it needs to show high rates of submissive behaviour towards the owner [29].

We hypothesized that serotonin and 5-HT_1A_ regulate aggressive and submissive behaviours of the focal ‘intruders’ as well as their winning likelihoods and contest durations in asymmetric competition tests, and that the neurotransmitter and its receptor will have similar effects on behaviour. In particular, we predicted that an increase of the serotonin concentration leads to a decrease of aggression [e.g. 18,34,35]. As the serotoninergic pathway influences aggression and submission in opposite directions [22], we further predicted that increased serotonin enhances submissive behaviour, and reduces contest duration. If this is the case this would mean that serotonin enhances social competence, as increased frequencies of submission are the appropriate response of *N. pulcher* in nature when they aim for acceptance by territory owners. When serotonin concentrations are lowered, we predicted the opposite effects on social behaviour (e.g. 36). Furthermore, a decrease in serotonin and a consequent increase in aggression may increase the probability of winning the contest. The activation of 5-HT_1A_ has been shown to affect aggression in a competitive context in a similar way than enhanced serotonin [14]. As we assume that the modulation of social behaviour through serotonin is mainly regulated by 5-HT_1A_, we predicted that the 5-HT_1A_ agonist will have similar effects than increasing serotonin concentration in the synaptic cleft, and the receptor antagonist to have similar effects than the serotonin-reducing drug.

## 3. Methods

### 3.1. Study Species

*N. pulcher* is a cooperatively breeding cichlid that is endemic to Lake Tanganyika, living in stable social groups with a size-based hierarchy. Groups consist of a dominant breeder pair and one to 20 subordinate individuals [6]. Subordinate individuals delay dispersal from the natal territory, often until adulthood. They help raising the dominant’s offspring by engaging in alloparental care, territory defence and maintenance as form of payment for being allowed to stay in the protection of the natal territory [36–39]. In *N. pulcher* severe predation risk is assumed to have selected for the species sociality [31,40], and therefore access to shelter places at a territory is crucial for their survival [6].

### Subjects and housing conditions

We used 160 fish to test the role of serotonin on the modulation of social behaviour. The fish were kept in their home tanks until testing. Eighty of those fish were assigned to be the focal fish, which received a drug treatment before testing and to take the role of an ‘intruder’ in an asymmetric competition test. They were paired with an individual matched by sex and size (± 0.05 cm; referred to as ‘owner’; N=80 owners). Forty focal fish were used in the serotonin enhancing experiment; they had a standard length (SL) of 3.57 ± 0.07 cm (mean ± se). Ten fish each received one of three dosages of Fluoxetine, and 10 fish received a control treatment. The 40 other focal fish were used in the serotonin-reducing experiment and had an SL of 3.97 ± 0.05 cm. Again, ten fish each received one of three dosages of PCPA and the remaining ten fish received a control solution. Sexes were balanced within treatments with five males and five females each per treatment; only in the Fluoxetine treatment with 10 µg/gbw six males and four females were tested.

To study the role of 5-HT_1A_ on the modulation of social behaviour we used another 40 fish, which were expose to a single dosage of receptor agonist, a single dosage of the receptor’s antagonist, and of saline as a control (details in drug applications). The fish were housed in four 100-L tanks for the duration of the study, which were equipped with three flower pot halves and semi-transparent plastic bottles and opaque plastic tubes on the water surface serving as shelters. The 40 fish were kept in groups of 10 fish per tank, separated by their assigned role (focal/non-focal fish) and by sex until testing. Twenty fish were assigned to be the focal fish and to take the role of the intruder during trials; they had an SL of 3.56 ± 0.04 cm. The sexes were balanced within treatments.

All experimental fish were bred and housed at the Ethological Station Hasli, Institute of Ecology and Evolution, University of Bern which is a licensed facility for cichlids (license number BE 4/11, Veterinary Office of the Kanton Bern). All experimental procedures were approved by the Veterinary Office of the Kanton Bern, licence 93/18, and were carried out in accordance with the standards of the National Research Council’s Guide for the Care and Use of Laboratory Animals and the EU directive 2010/63/EH for animal experiments. All tanks for holding the fish before and during experimental trials were equipped with a layer of sand and a biological filter. The water temperature was kept at 27 ± 1°C and the light:dark cycle was set to 13:11 hours, simulating natural light conditions in Lake Tanganyika [29].

### Experimental set-up

For each trial of the manipulation of either serotonin or the receptor a focal fish was temporarily transferred to one of four test tanks. These were 20-L tanks with opaque, removable PVC partitions dividing the tanks in two equally sized compartments. In the middle of one of the compartments half of a flowerpot was placed with the opening facing the opaque partition. Each of the compartments was equipped with an air stone to ensure sufficient aeration. The back and the two smaller sides of each test tank were covered with white plastic for reducing the disturbance of the fish. The compartment, in which the flowerpot half was placed, was balanced between left and right across tanks but it was kept at the same side for all trials of the same individual.

### Drugs

8-OH-DPAT (H8520), Way-100635 (W108), Fluoxetine (F132) and 4-Chloro-DL-phenylalanine (PCPA; C6506) were purchased from Sigma Aldrich (Deisenhofen, Germany). Drugs were dissolved in 0.9% NaCl to the respective concentration. Three concentrations were used for the manipulation of Fluoxetine (2.5, 5, and 10 µg/gbw) and of PCPA (2.5, 5, and 10 µg/gbw) based on previous fish studies [19,35,41,42]. In the 5-HT_1A_ manipulation, only one concentration of 8-OH-DPAT (1 μg/gbw) and WAY-100635 (1.5 μg/gbw) was used. The choice of this concentration is based on Stettler et al. [22], who had used three dosages of these drugs in our study species *N. pulcher* and had reported the strongest effects on the social behaviours of these fish with the concentrations used in the current study. In both experiments saline solution served as control. The drugs were stored at -25°C whenever they were not used.

### Drug applications

The Fluoxetine used to pharmacologically increase serotonin availability is a selective serotonin reuptake inhibitor (SSRI). The behavioural effects of SSRIs differ between acute and prolonged exposure to the drug. Acute SSRI exposure can lead to anxiogenic responses [43,44]. To avoid such acute, anxiogenic response in our experiment, we repeatedly administered fluoxetine in the food for 7 consecutive days. To do so, we produced agar-agar pellets of 0.50 cm height and 0.55 diameter supplemented with fish food containing thawed krill and either with solutions of one of the drugs in one of the concentrations or with saline solution. The amount of the drug solution added was standardized by the weight of the fish, so 3 μl/gbw were added to each pellet. To start the treatment (i.e., at the morning of day 1), a focal fish (‘intruder’) was placed in the compartment of a 20-L experimental tank without a shelter. At the same time, a stimulus fish was placed in the compartment with a shelter (‘owner’). The focal fish was fed the first pellet 2 h after placement in the tank, and then another pellet for the following 6 days always at the same time of day (12:00 hrs noon ± 1 h). The ‘owner’ received standard flake food at the same time. In one control trial of this experiment the owner managed to enter the compartment of the intruder on day 4. This trial was terminated and for this reason one trial of the control treatment is missing. In the mornings, before adding a new pellet, we removed any remainders of the pellets from the previous day and noted their approximate proportion of the original pellet size. Of 13.2 % of the pellets remainders were found. Twelve of the 40 focal fish left at least on one day the whole or parts of the pellet. It is thus possible that some focal fish did not get the full intended dosage. Following 7 days of feeding treatment, an asymmetric competition test was done in the morning of day 8 (see procedure below).

For the experiment where serotonin availability was reduced by a PCPA treatment, a focal fish (‘intruder’) and an ‘owner’ were placed in the two compartments of an experimental tank as described above. After one night of acclimatization, the focal fish was injected with either one of the three concentrations of the drug or the control saline solution in the morning of the next day (day 1). We performed intramuscular injections in the caudal muscle after the focal fish was anesthetized using KOI MED^®^ Sleep (Koi & Bonsai Zimmermann, Bühlertann, Germany). The amount was standardized respective to the body weight of the fish, namely 15 μl/gbw were injected [19,22,45]. The injections were done with a 0.5 ml insulin syringe. After the injection the focal fish was placed in a dark plastic bowl with an air stone for 5 min to recover from anaesthesia. Then it was placed back in its compartment. PCPA blocks tryptophan hydroxylase, which is responsible for the conversion of tryptophan to serotonin [7,46]. As serotonin excretion follows a circadian rhythm with serotonin depletion during the night [47], we waited 24 hours after the injection before doing the test in the morning of day 2 to make sure that the serotonin that was remaining in the system was used up at the moment of testing.

For the experiment investigating the role of 5-HT_1A_ agonist and antagonist, focal ‘intruder’ and ‘owner’ fish were placed in the experimental tanks as described above. After overnight acclimatization, the focal fish was injected after which the asymmetric competition trial started. Each focal fish was injected three times, once with the agonist, once with the antagonist, and once with saline solution, in a balanced treatment order. Between successive trials the fish received a break of at least 3 days for recovery and to ensure that the trials were independent. The anaesthesia, injection and recovery procedure were performed as described above for the PCPA manipulation. Thirty minutes after the injections, the asymmetric competition test was performed as described below.

### 3.2. Asymmetric competition test

Before the start of the asymmetric competition test each focal fish was video-recorded for 5 min with Sony Handycams mounted on tripods (type Sony HDR-PJ260 or Sony HDR-CX550) to get a recording of its spontaneous activity (i.e., when the fish was alone). Afterwards we started the asymmetric competition test by lifting the dividing partition, allowing the fish to interact. From the moment of lifting the divider, we video-recorded the trials for 20min, while the experimenter (PS) left the room to avoid distraction of the fish. After each trial all fish were transferred back to their respective holding tanks.

Afterwards the video recordings were analysed in real time using the software ‘Observer’, version 5.0.25 (Noldus, the Netherlands, 2003) by PS, who was blind to the treatment. Each asymmetric competition trial was analysed from contest start until the end of the contest over the shelter or for a maximum of 20 min (see 29,47). Contest start was defined by the moment the first contestant crossed the centre line of the tank. The end of the contest was defined by the time point at which a contest was decided by the intruder winning or losing the shelter (see below), or after a maximum of 20 min with no clear winner or loser (‘undecided’ contests). Undecided contests occurred as follows: in the fluoxetine experiment, 17 out of 39 trials; in the PCPA experiment, 14 out of 40 trials; in the 5-HT_1A_ experiment, 12 out 59 trials.

We scored all overt aggressive behaviours (bite, ram, mouth-fight), restrained aggression (fin spread, head down, opercula spread, approach) and submissive behaviours (tail quiver, escape/evade and hook display; for a detailed ethogram see Table S1). We also scored the time until the contest started, its duration and if the focal fish won or lost the contest. Winning was defined as (1) the focal intruder fish being tolerated by the owner within 3 cm of the shelter, or (2) the focal fish taking over the shelter entirely. Losing was defined as the intruder being evicted by the owner either staying far away from the shelter or being chased away from it, when trying to approach it. One video file of the experiment targeting the role of the receptor was corrupt, thus we lost one replication for the control treatment of the receptor experiment.

### Statistical analysis

All statistical analyses were done with the software R version 3.4.2 [49] with the interface R Studio version 1.1.383 (RStudio team, 2016) using the package ‘lme4’ [50]. The data of the experiments manipulating the serotonin availability were analysed using linear models (ANOVA) for activity and contest duration and negative binomial generalized linear models (GzLM) for overt aggression (bite + ram), restrained aggression (fin spread + head down + opercula spread) and submission (tail quiver + hook display; see Table S1 for a description of the behaviours). The data of the Fluoxetine trials and PCPA trials were analysed separately. Drug ID, sex, size, activity (activity included as covariate for all models except the one on activity) and a factor ‘outcome’ with three levels (focal fish either won, lost, or the contest was undecided) were included as fixed factors. The duration of the contest was included in the offset (except in the model when contest duration was the dependent variable. In the model analysing submission, owner aggression was included as fixed factor as aggression by a conspecific often directly elicits submission in *N. pulcher*. For the model on the contest duration only the data of trials where the contest was terminated before 20 min and thus had a clear winner and loser were analysed (Fluoxetine: N=23, PCPA: N=23).

The data of the experiment investigating the 5-HT_1A_ manipulation were analysed by linear mixed effect models (LMMs) for contest duration and activity. The frequency of overt aggression, restrained aggression and submissive displays (tail quiver, hook display) were analysed with generalized mixed effect models (GLMMs) following a Poisson distribution. The duration of the fight was included in the offset and the optimizer ‘bobyqa’ was used to control for model convergence. Treatment (DPAT, WAY and control), contest sequence (whether it was the first, second or third contest), sex, size, activity and contest outcome at the end of contest were included as fixed factors; (only in the model analysing treatment effects on activity as dependent variable, ‘activity’ was not included as fixed factor). Focal ID was included as a random factor to account for non-independence of the data. As above owner aggression was included as fixed factor in the model analysing submission, and for the model on the duration of the fight only the data of trials where the fight was decided before the maximal time of 20 min were used (N=41). We did a proportion test to compare the proportion of winners between the three treatments.

All models were simplified using AIC-based model selection, by performing a stepwise backward selection (Zuur et al. 2009) starting with the full model and always removing the variable that maximally reduced the AIC.

## 4. Results

### Manipulation of serotonin availability

#### Aggression

Restrained aggression was decreased by the two higher concentrations of Fluoxetine compared to the control treatment (Fluoxetine 5: P=0.038, Fluoxetine 10: P=0.007; nbGzLM, see Table 1a for full model results; Fig. 1a). The amount of overt aggression shown by the focal fish was not changed by Fluoxetine (Table 1b, Fig. 1b).

**Figure 1:**
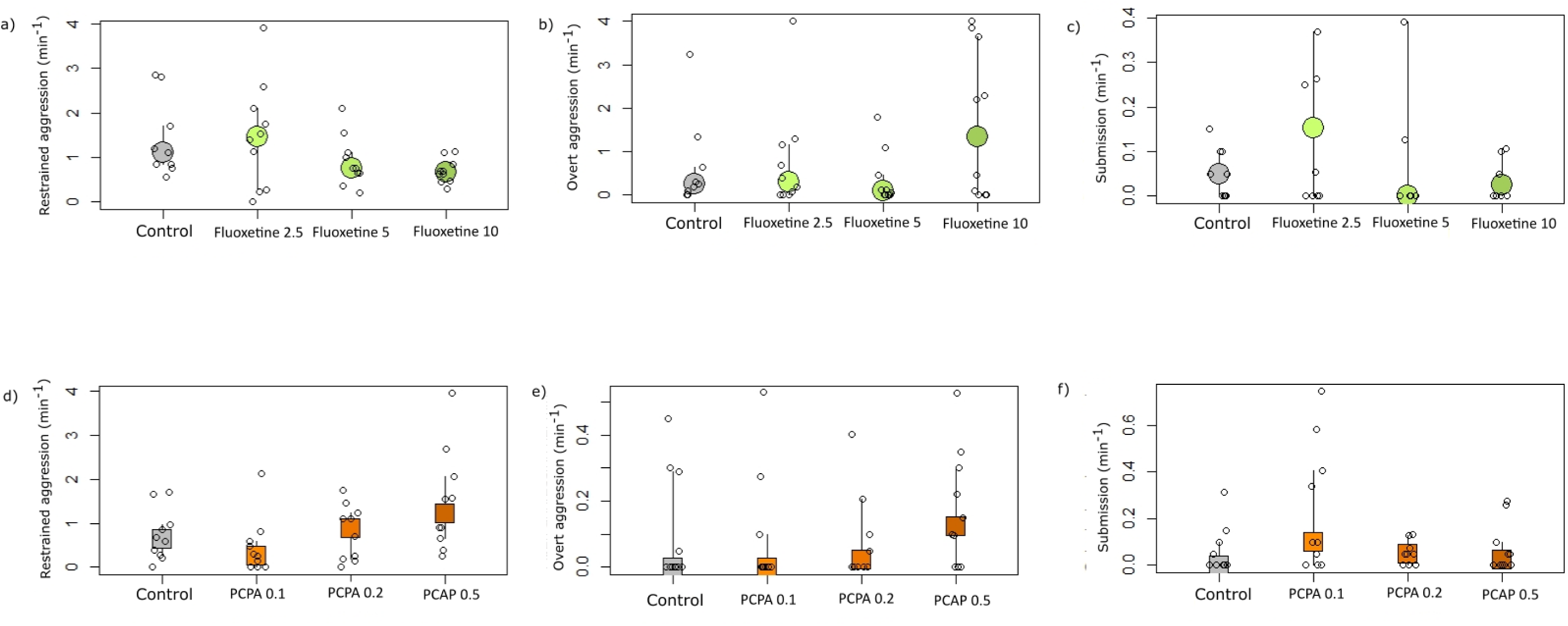
Manipulation of serotonin availability. The effects of different Fluoxetine dosages on (a) restrained aggression; (b) overt aggression and (c) submission and the effects of PCPA dosages on (d) restrained aggression; (e) overt aggression; (f) submission. Medians and interquartile ranges are shown.

**Table 1:** The results of the negative binomial GzLMs on the effect of Fluoxetine on (a) restrained and (b) overt aggression, (c) submissive behaviour, and the ANOVAs on (d) activity and (e) contest duration. (a-d: N=39, e: N=23).

| Factors | Estimate ± SE | z/t | p-value |
| --- | --- | --- | --- |
| **a) Restrained aggression** |  |  |  |
| (Intercept) | -3.73864 ± 0.17895 | -20.89 | **<0.0001** |
| Fluoxetine 2.5 | 0.02047 ± 0.25583 | 0.080 | 0.94 |
| Fluoxetine 5 | -0.53964 ± 0.26045 | -2.072 | **0.038** |
| Fluoxetine 10 | -0.69269 ± 0.25716 | -2.694 | **0.0071** |
| **b) Overt aggression** |  |  |  |
| (Intercept) | -10.6719 ± 1.9651 | -5.431 | **<0.0001** |
| Fluoxetine 2.5 | -0.9208 ± 0.9647 | -0.954 | 0.34 |
| Fluoxetine 5 | 0.1934 ± 0.6868 | 0.282 | 0.78 |
| Fluoxetine 10 | 0.8121 ± 0.6593 | 1.232 | 0.22 |
| Size | 1.3656 ± 0.5107 | 2.674 | **0.0075** |
| Outcome: undecided | 1.5688 ± 0.5462 | 2.872 | **0.0041** |
| Outcome: winner | 2.8236 ± 1.0204 | 2.767 | **0.0057** |
| **c) Submission** |  |  |  |
| (Intercept) | -6.0668 ± 0.8062 | -7.525 | **<0.0001** |
| Fluoxetine 2.5 | 1.7132 ± 1.0441 | 1.641 | 0.10 |
| Fluoxetine 5 | 1.0386 ± 0.8830 | 1.176 | 0.24 |
| Fluoxetine 10 | 0.8473 ± 0.8472 | 1.000 | 0.32 |
| Outcome: undecided | -1.0286 ± 0.6565 | -1.567 | 0.12 |
| Outcome: winner | -2.4013 ± 1.1729 | -2.047 | **0.041** |
| **d) Activity** |  |  |  |
| (Intercept) | 0.588889 ± 0.066860 | 8.808<sup>t</sup> | **<0.0001** |
| Fluoxetine 2.5 | 0.001111 ± 0.092160 | 0.012<sup>t</sup> | 0.99 |
| Fluoxetine 5 | -0.078889 ± 0.092160 | -0.856<sup>t</sup> | 0.40 |
| Fluoxetine 10 | -0.113889 ± 0.092160 | -1.236<sup>t</sup> | 0.225 |
| **e) Duration of contest** |  |  |  |
| (Intercept) | -517.10 ± 467.12 | -1.107<sup>t</sup> | 0.283 |
| Fluoxetine 2.5 | -377.02 ± 257.84 | -1.462<sup>t</sup> | 0.16 |
| Fluoxetine 5 | -116.58 ± 240.95 | -0.484<sup>t</sup> | 0.63 |
| Fluoxetine 10 | -11.92 ± 250.57 | -0.048<sup>t</sup> | 0.96 |
| Size | 337.72 ± 126.84 | 2.663<sup>t</sup> | **0.016** |
| Outcome: winner | 423.32 ± 196.94 | 2.150<sup>t</sup> | **0.046** |

The lowest concentration of PCPA significantly decreased restrained aggression compared to the control (P=0.044; nbGzLM, Table 2a, Fig. 1d). The higher two concentrations had no effect. Overt aggression was not influenced by either of the concentrations of PCPA (Table 2b, Fig. 1e).

**Table 2:**
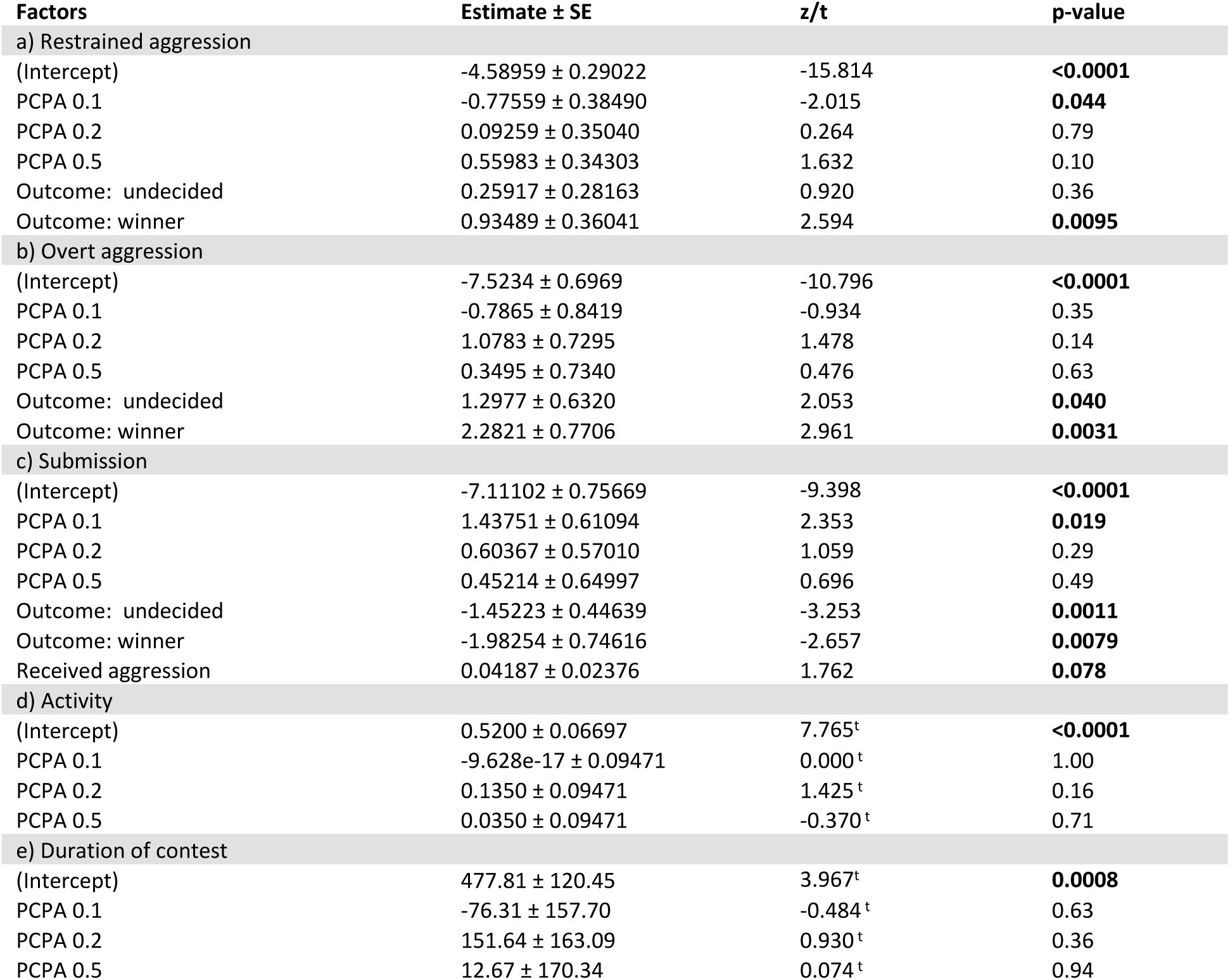
results of the negative binomial GzLMs on the effect of PCPA on (a) restrained and (b) and overt aggression, (c) submission, and the ANOVAs on (d) activity and (e) contest duration. (a-d: N=40, e: N=23).

#### Submission

None of the concentrations of Fluoxetine significantly influenced submissive behaviour (Table 1c, Fig. 1c). The lowest concentration of PCPA led to an increase of submissive behaviour shown by the focal fish compared to the control (P=0.019; nbGzLM, Table 2c, Fig. 1f). The other two concentrations had no effect.

#### Activity

None of the three concentrations of Fluoxetine influenced the activity of the focal fish (Table 1d). Likewise, the activity was not influenced by any of the concentrations of PCPA (Table 2d).

#### Contest duration

The duration of the contest was not influenced by either of the three concentrations of Fluoxetine (Table 1e) or PCPA (Table 2e).

#### Winning

The proportion of focal fish being the winner differed in the Fluoxetine experiment across drug dosages, both when calculated for decided contests only (winners, losers, Proportion test, df=3, χ^2^=9.94, p=0.019, n=23) or for all contests (winners, losers, undecided contests, Proportion test, df=3, χ^2^=16.63, p<0.001, n=39). Visual inspection of Fig. S1 suggests that these differences occur because after application of the lowest Fluoxetine concentration (but not other fluoxetine concentrations neither from the control solution) several focal individuals ended the contest as winners. In the experiment with PCPA, the proportion of winners did not differ between the treatments both when calculated for decided contests only (Proportion test, df=3, χ^2^=1.92, p=0.59, n=26) or when calculated for all contests (Proportion test, df=3, χ^2^=1.90, p=0.59, n=40).

### Agonist and Antagonist of the serotonin 1A receptor

#### Aggression

Application of the agonist of 5-HT_1A_, 8-OH-DPAT, significantly reduced restrained aggression compared to the control (Table 3a, Fig. 3a), but did not affect overt aggression (Table 3b). The receptor’s antagonist, Way-100635, did not affect restrained or overt aggression (Table 3a, Fig. 3a). There was also an effect of trial order: in the second and the third contest, focal fish showed less restrained and overt aggression than in the first contest (Table 3a,b). Individuals that were classified as winners of a contest showed more restrained and overt aggression during the contest than losers (Table 3a,b).

**Tab 3:** Results of the GLMMs on the effects of the 5-HT_1A_ manipulation on (a) restrained aggression, (b) overt aggression, and (c) submission, and the LMMs on (d) activity and (e) the contest duration. (a-d: N=59, e: N=41)

| Factors | Estimate ± SE | z/t | p-value |
| --- | --- | --- | --- |
| **a) Restrained aggression** |  |  |  |
| (Intercept) | -5.5966 ± 0.3415 | -16.387 | **<0.0001** |
| 8-OH-DPAT | -1.0611 ± 0.3669 | -2.892 | **0.0038** |
| WAY-100635 | 0.1435 ± 0.1656 | 0.867 | 0.386 |
| Activity | 1.3385 ± 0.5666 | 2.363 | **0.0181** |
| Sequence 2 | -0.4302 ± 0.1886 | -2.281 | **0.0226** |
| Sequence 3 | -1.1723 ± 0.2434 | -4.817 | **<0.0001** |
| Outcome: undecided | 0.3752 ± 0.2893 | 1.297 | 0.195 |
| Outcome: winner | 1.0753 ± 0.3995 | 2.691 | **0.0071** |
| **b) Overt aggression** |  |  |  |
| (Intercept) | -9.0292 ± 0.7084 | 12.746 | **<0.0001** |
| 8-OH-DPAT | -1.2158 ± 0.9017 | -1.348 | 0.1775 |
| WAY-100635 | -0.1864 ± 0.4923 | -0.379 | 0.7049 |
| Activity | 1.6458 ± 1.4075 | 1.169 | 0.2423 |
| Sex: male | 2.0702 ± 0.5824 | 3.554 | **0.0004** |
| Sequence 2 | -0.2876 ± 0.5367 | -0.536 | 0.5921 |
| Sequence 3 | -1.2729 ± 0.5373 | -2.369 | **0.0178** |
| Outcome: undecided | 0.7738 ± 0.6449 | 1.200 | 0.2302 |
| Outcome: winner | 2.0087 ± 0.7026 | 2.859 | **0.0042** |
| **c) Submission** |  |  |  |
| (Intercept) | -5.9593 ± 0.4214 | -13.99 | **<0.0001** |
| 8-OH-DPAT 1 | 0.0352 ± 0.3666 | 0.096 | 0.92 |
| Way-100635 1.5 | -0.1666 ± 0.6393 | -0.631 | 0.51 |
| Received aggression | 0.0140 ± 0.0063 | 2.206 | **0.027** |
| Outcome: undecided | -0.4698 ± 0.405 | -1.159 | 0.24 |
| Outcome: winner | -17.65 ± 1956 | -0.009 | 0.98 |
| **d) Activity** |  |  |  |
| (Intercept) | 0.38350 ± 0.04746 | 8.081<sup>t</sup> | **<0.0001** |
| 8-OH-DPAT 1 | -0.23100 ± 0.05633 | -4.101<sup>t</sup> | **0.00021** |
| Way-100635 1.5 | 0.09900 ± 0.05633 | 1.757<sup>t</sup> | 0.087 |
| **e) Duration of contest** |  |  |  |
| (Intercept) | 525.86 ± 66.97 | 7.852<sup>t</sup> | **<0.0001** |
| 8-OH-DPAT 1 | -37.98 ± 61.73 | -0.615<sup>t</sup> | 0.54 |
| Way-100635 1.5 | 85.46 ± 68.87 | 1.241<sup>t</sup> | 0.22 |
| Sequence 2 | -243.65 ± 59.71 | -4.080<sup>t</sup> | **0.00025** |
| Sequence 3 | -285.70 ± 64.72 | -4.414<sup>t</sup> | **<0.0001** |
| Sex | -139.80 ± 51.63 | -2.708<sup>t</sup> | **0.010** |

**Figure 2:**
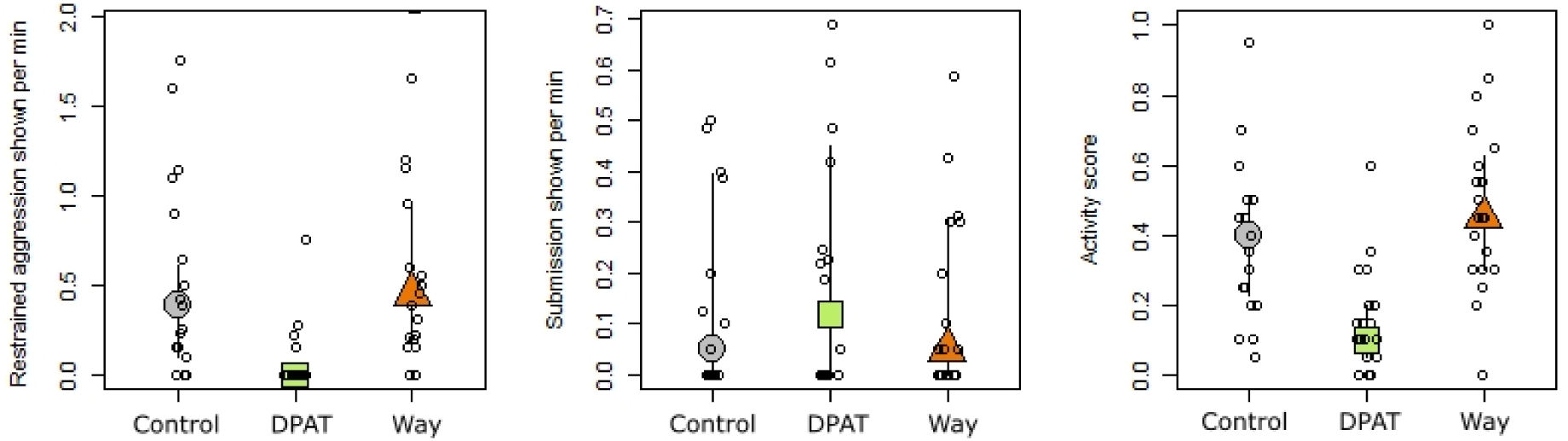
Effects of the serotonin 1a receptor manipulation on: (a) frequency of aggression per minute of the contest, (b) frequency of submission per minute of the contest, (c) activity. Plots show medians and interquartile ranges, small circles represent the individuals. Grey circles: medians of control treatment; green squares: medians of DPAT treatment; orange triangles: medians of WAY treatment. Medians and interquartile ranges are shown.

#### Submission

Neither of the two drugs caused a change in submissive behaviour compared to the control (Table 3c, Fig. 3b). The submissive behaviour increased when focal fish received more aggression (Table 3c).

#### Activity

With 8-OH-DPAT the individuals were significantly less active than with saline solution (P<0.001; LMM, Table 3d, Fig. 3c). Way-100635 tended to have the opposite effect, namely leading to higher activity compared to the control (P=0.087; LMM, Table 3d, Fig. 3c).

#### Contest duration

None of the drugs had a significant influence on the duration of the contest (Table 3e). The second and the third contest of a pair were significantly shorter than the first (Table 3e).

#### Winning

The proportion of winners did not differ between the three treatments. There was no difference in the proportions of winners neither when calculated only for decided contests (winners, losers, P=0.35; Proportion test, df= 2, χ^2^=2.21, n=47) nor when calculated over all contests (winners, losers, undecided contests, P=0.58; Proportion test, df= 2, χ^2^=1.10, n=59).

## 5. Discussion

Here we showed that serotonin modulates social competence in *N. pulcher*: the lowest concentration of PCPA increased submission relative to the aggression received by an opponent. However, this modulation does not occur via the activity of the receptor 5-HT_1A_. Both the serotonin-enhancing (Fluoxetine) and the serotonin-reducing (PCPA) drugs decreased restrained aggression, without affecting the expression of overt aggressive behaviour. As we predicted, pharmacological activation of the 5-HT_1A_ receptor decreased aggression in a competitive context, but it affected only restrained aggression.

### Manipulation of serotonin availability

Manipulation of serotonin availability influenced social competence by modulating aggressive behaviour and submissive behaviour relative to received aggression by the opponent during an asymmetric contest over a resource. Fluoxetine decreased restrained aggression at the higher two concentrations and increased the likelihood to win a contest at the lowest fluoxetine dosage. These results suggest that increased serotonin availability might enhance the likelihood to achieve a dominant status. PCPA decreased restrained aggression and increased submission at the lowest concentration, suggesting that restrained aggression and submission are inversely linked. The relationship between serotonergic activity and social status has been previously shown in rainbow trout [51] and other fish species (reviewed in [46]), but also in reptiles [10] and rodents [52]. As all dyadic aggressive interactions are stressful for the involved individuals [51], this leads to an increase in serotonergic activity and activation of the stress response system [46,53]. In teleost fish, after establishing a hierarchy, dominant individuals reduced their serotonergic activity while subordinate individuals kept their serotonergic activity elevated, shown by the higher concentration of the 5-HT metabolite, 5-hydroxyindoleacetic acid (5-HIAA; [46]). The link between social status and winning likelihood is not well understood. In *N. pulcher*, assigned rank does not seem to influence the likelihood to win a contest [54]. However, social rank influences restrained aggression, with dominant individuals showing less restrained aggression [54]. Restrained aggression is a form of aggressive behaviour that can be shown at the beginning of a contest, when the opponents are evaluating each other’s abilities [55]. In zebrafish, winning a fight caused a significant increase of brain serotonin, while losers showed an increase of the serotonin metabolite 5-HIAA in the hypothalamus [56]. Taking into account the behavioural results observed from our 5-HT availability manipulation, we hypothesize that (i) a pharmacological increase of 5-HT with fluoxetine induced a “dominant-like” state which caused the reduction in restrained aggression and an increased likelihood to win a contest; (ii) pharmacological decrease of 5-HT availability induced a “subordinate-like” state, resulting in reduced restrained aggression as well, but also increased submission per received aggression. Our results therefore highlight the relative importance of 5-HT in determining social status.

Regarding the dosage-depend effects on behaviour, we expected that there would be dosage-dependent behavioural effects, with higher dosages of fluoxetine and PCPA having more pronounced effects [22,45,46]. We further expected that the administered drugs would induce opposite behavioural effects. Interestingly, however, both drugs caused similar effects on restrained aggression, at the intermediate and highest concentrations of fluoxetine (Fig. 1a), and at the lowest concentration of PCPA (Fig. 1d). These results might be explained by partial blockage of serotonin synthesis by a low concentration of PCPA, which may lead to an increase of serotonergic activity, regulated over a negative feedback loop regulated by postsynaptic 5-HT_1A_ [27]. The same may apply for Fluoxetine for which it is also possible that a partial blocking of the serotonin transporters leads to an actual decrease of serotonin availability. Chronic administration of SSRIs can lead to desensitisation of somatodendritic 5-HT1a autoreceptors, which consequently results in reduced ability to serotonergic transmission [57]. We hypothesize that our repeated administration of the higher dosages of fluoxetine might have induced a desensitization of the 5-HT_1A_ autoreceptors, which impaired the ability of the fish to release serotonin to the synaptic cleft. Further experiments are necessary to validate this hypothesis and to deepen our knowledge on the mechanistic basis of PCPA and fluoxetine in regulating social competence. Nonetheless, our concentration-specific results show the importance of performing a dosage-dependent study by testing the effects of multiple dosages of drugs when first investigating it in a particular species. Another reason for the different effects between the drugs could be the different administration methods. For PCPA we analysed an acute response, while fluoxetine was repeatedly administered for one week via food.

### Serotonin 1A receptor manipulation

The 5-HT_1A_ pharmacological activation significantly decreased restrained aggression and did not have an effect on submissive behaviour, while the pharmacological blockage of the receptor did not have an effect on any of the analysed behaviours during the contests. Regarding the overall activity of the focal fish, 5-HT_1A_ activation decreased activity and blockage tended to increased it (Table 3d). To avoid the influence of activity on the analysis of social behaviours, we statistically controlled for it in our models (see Methods: Statistical analysis). We showed that the serotonin 1A receptor does play a role in the regulation of restrained aggression in a competitive context. However, we do not have evidence for a role of the receptor’s activity in the modulation of submissive behaviour, possibly due to the activity of other serotonergic receptors or other neurophysiological systems, such as dopamine [45]. Like in our study, in *Betta splendens*, the 5-HT_1A_ agonist decreased restrained aggression (i.e. opercular spreads), while the antagonist of the receptor did not influence restrained aggression [14]. We have recently shown in *N. pulcher* that environmental context has to be taken into account when analysing the role of neurophysiological systems in regulating social behaviour, as when individuals are exposed to different environmental challenges the social demands from other group members also change [45]. For instance, when belonging to a social group, 5-HT_1A_ activation increased total aggression among group members of *N. pulcher* [22], whereas in this study aggression was decreased by the same manipulation. Such discrepancy between results can be explained by the differences in the environmental contexts, which require different responses, highlighting the importance of both social and environmental contexts when studying the neuronal mechanisms of behaviour. Regarding the lack of an effect on submission, there are a number of other serotonergic receptors beside the serotonin 1A receptor, which play a role in the modulation of social behaviour, for example the serotonin 2A and the serotonin 7 receptor [14,20,21]. It is thus possible that one of those receptors which we did not manipulate is responsible for regulating the effects of serotonin on submission.

### General discussion

Our results suggest a role of serotonin in the modulation of aggressive and submissive behaviour in *N. pulcher*. Within the families, *N. pulcher* maintain a linear social hierarchy, which is stabilized by showing aggressive behaviour when being dominant over an individual and responding by submissive behaviour to received aggression when being subordinate [6]. As we show, serotonin modulates these social behaviours, revealing the relative importance of the serotonergic system in regulating social competence. Social competence is particularly important for subordinate individuals, as their permanence in families and access to the protection of a territory and, consequently, their survival is dependent on showing the appropriate behaviour towards dominants [29,58,59]. Our results elucidate the role of the serotonergic system in regulating social competence, and generally highlight the importance of social and environmental contexts in behavioural neuroscience.

## 6. Acknowledgements

We thank Svante Winberg for his helpful comments while designing this experiment and to an earlier version of this manuscript, and Evi Zwygart for logistical support. We acknowledge financial support by the Swiss National Science Foundation (SNSF, project 31003A_179208 to BT).

## 7. Authors contribution

Conceptualization: D.F. Antunes, P. Stettler, B. Taborsky; methodology: D.F. Antunes, P. Stettler, B. Taborsky; data collection: P. Stettler; Analysis: D.F. Antunes, P. Stettler, B. Taborsky; writing/editing: D.F. Antunes, P. Stettler, B. Taborsky. Funding: B. Taborsky.

