## Supplement for "The role of serotonin in modulating social competence in a cooperatively breeding fish"

Content: Supplementary Table S1, supplementary Figure S1

Table S1: Ethogram of *Neolamprologus pulcher* used in this study

| Behavioural category | Behaviour | Description |
|---|---|---|
| Restrained aggression | Fin spread | All fins are maximally spread, the body is kept in a stiff posture |
| | Head down | Like fin spread, but the body is tilted forwards |
| | Opercula spread | Spreading of the operculum and lowering of the branchiostegal membrane. Often occurring while the fish is performing a fast linear approach towards the opponent, stopping shortly before physical contact. |
| | Approach/chasing | Fast approach towards another fish which is followed by evade or escape of the opponent |
| Overt aggression | Ram | Fast linear approach towards another fish ending with physical contact, mouth closed |
| | Bite | Biting another fish |
| Submission | Tail quiver | Caudal peduncle, tail fin and back end of dorsal fin are intensively vibrating while the unpaired fins are folded |
| | Hook display | Fast swimming to and away from another fish (bow swimming) with a light touch at the apex of the bow. |
| | Evade | Fish makes a small movement away from an approaching fish |
| | Escape | Fish moves away as fast as possible, fleeing from an attacking fish |

Figure S1: Contest outcome after fluoxetine administration. Number of contests where the focal fish was losing, undecided contests, and number of contests with the focal fish winning per treatment.

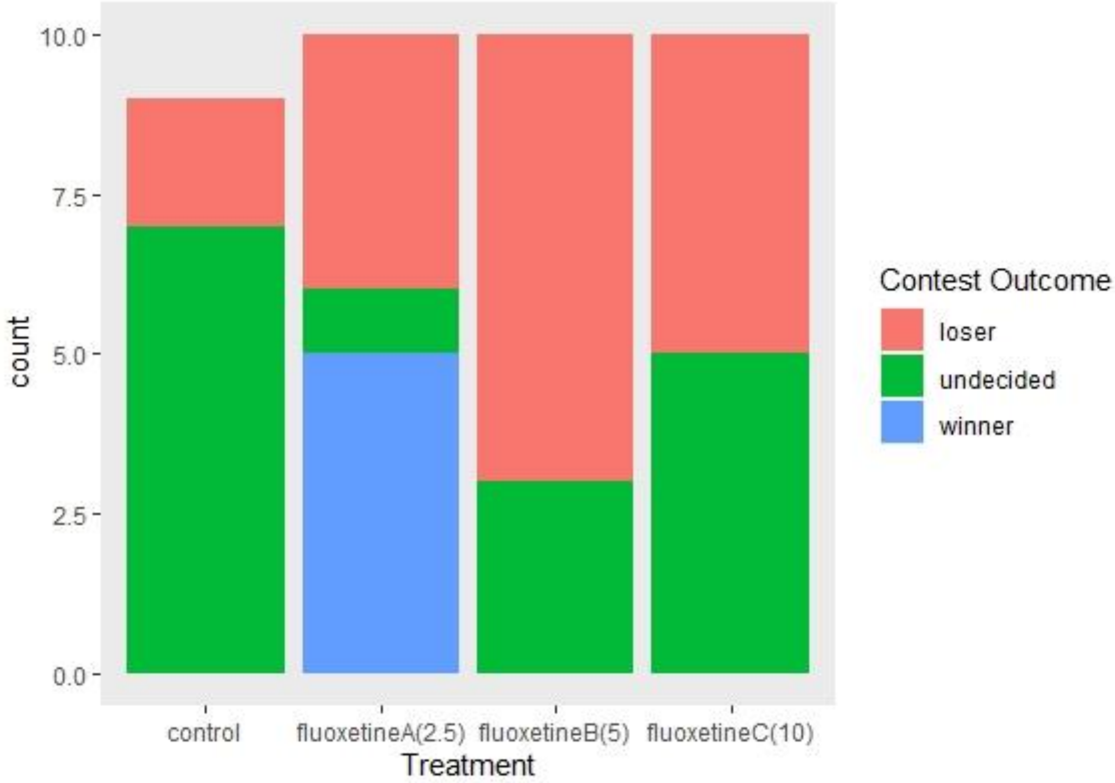
